# A biphasic trajectory for maize stalk mechanics shaped by genetic, environmental, and biotic factors

**DOI:** 10.1101/2025.03.03.641235

**Authors:** Irene I. Ikiriko, Ashley N. Hostetler, Jonathan W. Reneau, Alyssa K. Betts, Erin E. Sparks

**Affiliations:** Department of Plant and Soil Sciences, University of Delaware, Newark, DE USA; Division of Plant Science and Technology, University of Missouri, Columbia, MO USA; Department of Plant and Soil Sciences, University of Delaware, Georgetown, DE USA; Donald Danforth Plant Science Center, St. Louis, MO USA

**Keywords:** Stalk flexural stiffness, bending modulus, 3-point bend, Pythium root rot

## Abstract

Stalk mechanical properties impact plant stability and interactions with pathogenic microorganisms. The evaluation of stalk mechanics has focused primarily on the end-of-season outcomes and defined differences among inbred and hybrid maize genotypes. However, there is a gap in understanding how these different end-of-season outcomes are achieved. This study measured stalk flexural stiffness in maize inbred genotypes across multiple environments and in maize commercial hybrid genotypes under different disease states. Under all conditions, stalk flexural stiffness followed a biphasic trajectory, characterized by a linear increase phase and a sustained phase. Within a genotype, the environment or disease state altered the rate of increase in the linear phase but did not impact the timing of transition to the sustained phase. Whereas between genotypes, the timing of transition between phases varied. Destructive 3-point bend tests of inbred stalks showed that the trajectory of stalk mechanics is defined by the bending modulus, not the geometry. Together, these results define a biphasic trajectory of maize stalk mechanics that can be modulated by internal and external factors. This work provides a foundation for breeding programs to make informed decisions when selecting for optimized stalk mechanical trajectories, which is necessary for enhancing resilience to environmental stresses.

**HIGHLIGHTS:**

- A common trajectory of stalk mechanics was found across genotypes and conditions.
- Transition between phases of stalk mechanics is inherent to genotype.
- The rate of linear increase phase is modulated within a genotype by environmental or biotic factors.
- Material properties determine the trajectory of stalk mechanics.

## INTRODUCTION

Maize (*Zea mays*) is an agricultural staple crop cultivated globally for food, biofuel, and livestock feed. Maize production is dependent on the generation of commercial hybrid germplasm that is resilient to growth in different environments. These hybrids are generated by selecting desirable traits in inbred genotypes which are then crossed. Inbred genotypes are typically used in research studies to link genotype to phenotype, however, desirable traits identified within inbred genotypes do not necessarily translate to their hybrid offspring. This emphasizes the need to evaluate both research inbred and commercial hybrid germplasm. For all germplasm, core aspects of plant production are plant stability and pathogen-interactions, both of which are determined by biomechanics.

Biomechanics describes the relationship between force and displacement in living systems with displacements applied in different contexts. Plant mechanics can be divided into two sequential mechanical modes of deformation: elastic and plastic (Niklas and Spatz, 2012). Elastic deformation is characterized by non-permanent changes and is reported as stiffness. Plastic deformation is characterized by permanent changes and is reported as strength. Both stiffness and strength are used to quantify the mechanical behavior of plants. In maize, the stalk is the structural axis that supports the above-ground organs, including the grain-producing ear. Thus, stalk mechanics is important for crop production and one factor that influences resistance to lodging (i.e., mechanical failure) and interaction with microorganisms.

Several studies have quantified maize stalk mechanics (both stiffness and strength) at a single time point and shown variation at the end-of-season for both inbred and hybrid genotypes (Kunduru et al., 2023; Oduntan et al., 2024; Sekhon et al., 2020; Xue et al., 2020; Zhang et al., 2018). Achieving these end-of-season differences must be the outcome of different trajectories of stalk mechanics during plant growth. One study defined part of this trajectory by comparing stalk stiffness at two growth stages, and showed an increase from the late vegetative to the late reproductive stage (Reneau et al., 2020). However, the overall trajectory of stalk mechanics to achieve this increase is unknown. One possible trajectory could be a constant increase of stalk mechanics throughout plant growth. Alternatively, there could be multiple phases, e.g., a phase of rapid increase followed by a phase of sustained or reduced increase. In destructive measurements of wheat, the stalk mechanics show a biphasic trajectory consisting of a linear increase followed by a sustained phase (Crook et al., 1994). It has not been reported which of these trajectories maize stalk mechanics follows nor whether these trajectories are specific to genotypes or can be altered by environmental or biotic factors.

Environmental factors such as varying the timing and rate of fertilizer application can impact stalk mechanics. In maize, reduction in nitrogen application increased stalk strength and lignin content, a critical secondary cell wall polymer (Ahmad et al., 2023). This outcome is consistent with results from other crops such as rice and wheat, which consistently show a reduction in fertilizer application resulted in increased stalk strength (Crook and Ennos, 1995; Zhang et al., 2017, 2014). Plant disease also influences stalk mechanics, since pathogens compromise the integrity of plant tissues or cause remobilization of nutrients (Xue et al., 2021). Many foliar and stalk rot diseases are not noted until later in the season and qualitative push or pinch tests can be used to indirectly evaluate mechanics (Jackson-Ziems et al., 2014). Pythium Root Rot (PRR), caused by soil-borne oomycetes within genera *Pythium*, *Elongisporangium*, and *Globisporangium* (Nguyen et al., 2022), is one disease that threatens maize production beginning from early season (Bickel and Koehler, 2021). While the most common outcome of *Pythium* infection is pre- or post-emergence seedling death, some plants survive the initial infection but yield poorly compared to healthy plants (Henrickson, M.G., Bickel, J.T., Betts, A.K., 2025). Plants that survive initial *Pythium* infection likely have altered mechanical trajectories, but this has not been defined. Understanding how environmental and biotic factors shape stalk mechanical properties requires quantitative measurements of stalk mechanics.

Maize stalk mechanics have been measured in the field using devices that treat the plant as a cantilever beam and quantify stalk flexural stiffness (*EI*) (Shah et al., 2017; Stubbs et al., 2022b). Field-based approaches are advantageous in that they provide in-situ quantification of mechanics. However, they cannot separate the contribution of plant geometry (second moment of area, *I,* which determines how forces are distributed) from the material properties (bending modulus, *E*) of the plant. In contrast, laboratory-based approaches, including 3-point bend tests, provide ex-situ quantification of mechanics that enables the separation of contributions from geometry and material properties (Niklas and Spatz, 2012). Yet, these approaches are also limited in that they only quantify a portion of the stalk, losing the contexts from the whole-plant and field environment. Combining field and laboratory approaches to measure stalk mechanics provides the most comprehensive assessment.

This study used a field-based device to quantify stalk flexural stiffness trajectories in maize inbred genotypes under different environments, and commercial hybrid genotypes that were symptomless and those that survived following early season symptoms of PRR. In all cases, there was a biphasic trajectory, a rapid increase phase followed by the transition to a sustained phase. Within a genotype, the rate of increase was modulated by environment and disease state, but the timing of transition was constant. Between genotypes, the timing of transition was variable.

Laboratory-based 3-point bend measurements showed that the trajectory of stalk flexural stiffness is driven by changes in material properties rather than geometry. Collectively, this work demonstrates a common trajectory of maize stalk flexural stiffness under different conditions but demonstrates that the shape of the trajectory can be modulated for plant resilience.

## MATERIALS AND METHODS

### Plant Material and Field Conditions

Seeds from the CIMMYT tropical inbred genotype CML258 and from the temperate inbred genotypes B73 and Mo17 were planted in 2021, 2023, and 2024. CML258 was planted on 20 May 2021; CML258, B73, and Mo17 were planted on 17 May 2023 and 20 May 2024. Across all years, seeds were planted in 2-6 replicate 3.66 m plots in Newark, DE USA. Fields were irrigated with an overhead linear irrigation system and were treated with pre-emergence herbicide (Lexar at 8.18 L ha^-1^ and Simazine at 2.81 L ha^-1^) and post-emergence herbicide (Accent at 0.05 L ha^-1^).

Soil insecticide (COUNTER 20 G at 6.16 kg ha^-1^) and fertilizer (ammonium sulfate 21-0-0 at 100.88 kg ha^-1^) were applied at planting, and at approximately one month after planting, the fields were side-dressed with 30% urea ammonium nitrate (at 374.16 L ha^-1^). In 2023, the field was not side-dressed due to being too wet for the tractor. Weather data for the Newark, DE field site can be found on the Delaware Environmental Observing System (http://www.deos.udel.edu) by selecting the Newark DE-Ag Farm station. Additionally, cumulative growing degree (GDD) data was downloaded from the NEWA growing degree day calculator (https://newa.cornell.edu/degree-day-calculator/) with Base 50°F parameters for our Newark Ag Farm location to compare data across years (**Figure S1A**). GDD is a calculation of heat units that can serve as a proxy for developmental progression of an individual genotype, since developmental transitions within a genotype will occur at set GDD.

Seeds from commercial hybrid genotypes were grown in production fields in Seaford, DE USA. Field 1 was planted with Axis 63H27 (Axis Seed) on 20 April 2023, Field 2 was planted with Channel 210-46 (Bayer Crop Science) on 19 April 2023, and Field 3 was planted with Axis 60W25 (Axis Seed) on 18 April 2023. All fields were planted at a population of 84,016 seeds per hectare (34,000 seeds per acre). Fields were scouted during the vegetative growth stage (V2–V3), for seedlings with stunting, chlorotic and necrotic tissue, or damping off to indicate presence of PRR. A subset of samples were collected and brought to the lab for isolate recovery and pathogen identification to verify *Pythium* presence within each field. Grids were then established in field areas showing symptoms of PRR by marking five rows spaced 76 cm apart. Rows were 5.3 m in length to simulate 4 square m (1/1000th of an acre). Four grids were established across the three fields. PRR symptomatic plants were identified and marked with a stake within each grid. Plants with severe symptoms were excluded to allow for repeated season long observations. A marked “pair” was completed by staking a symptomless corn plant immediately beside the symptomatic plant. Each grid contained five plant pairs (n=20 pairs). Pairs were evaluated throughout the growing season. Precipitation and weather data was recorded monthly by accessing the Delaware Environmental Observing System (http://www.deos.udel.edu) and selecting the Seaford, DE-Oak Grove Farm station.

### Non-Destructive Quantification of Stalk Mechanics

#### Device Design & Operation

A device, called the Pusher, was developed to non-destructively measure stalk flexural stiffness of large grain crops in the field. The Pusher was developed to improve data collection and precision compared to the previously described DARLING device <u>(Cook et al. 2019)</u>. While the DARLING has proven to be useful in collecting non-destructive stalk biomechanics data <u>(Reneau et al. 2020;</u> <u>Cook et al. 2019)</u>, it is manually operated and therefore subject to inherent human error. In contrast, the Pusher automates data collection by activating a leadscrew (described below) which reduces the variability and eliminates user-introduced error.

The Pusher device consists of a frame, powertrain, sensors, electronics, and software (**Figure S2**). Two different frames were used: one built from 3.81 cm aluminum extrusion square stock, and a lighter frame built from 2.54 cm aluminum extrusion square stock. Regardless of the frame being used, the powertrain was attached to a vertically adjustable plate (set at 0.54 m for the measurements in this study). The powertrain consists of a linear guide rail with a leadscrew driven carriage, brushed DC motor, and rechargeable battery. A 294 N (30 kg) load cell was attached to the carriage with an optional 3D printed attachment to center the stalk on the load cell. The internal electronics were run via an Arduino microcontroller connecting the motor driver, a load cell amplifier, and using the serial terminal to connect to custom Python software.

To acquire data, the load cell was placed in contact with the stalk. The device was stabilized by the user standing on the base of the frame. The “Start Test” button was pressed on the software, which activated extension of the leadscrew at 0.5387 cm/s for a total push length of 4.445 cm (**Video S1**). The device recorded data in approximately 0.05 cm increments. Displacement (cm) and force (N) were recorded with the device.

#### Data Processing

A linear regression was run with displacement as the independent variable and force as the dependent variable. The stalk flexural stiffness (*EI*) was calculated via the following equation:

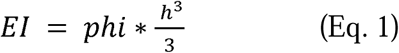

where phi is the slope of the linear regression (N/m) and h is the height of the applied displacement (m) (set at 0.54 m in this study).

#### Evaluation of Stalk Flexural Stiffness

The stalk flexural stiffness was quantified approximately weekly (weather permitting) for the duration of the experiment, beginning between 56-64 dap, ending between 132-141 dap for inbreds and beginning between 75-77 dap, ending between 130-132 dap for hybrids. In 2021, the stalk flexural stiffness was quantified for CML258 weekly in 40 plants. A sample size analysis <u>(Faul *et al*., 2007)</u> was run with this data and we determined that 4-6 plants were required to meet a statistical power of 0.05. We ensured this sample size was met by sampling 4-6 plants per plot for inbred lines. For hybrid lines, we started with 5 pairs per grid, but lost samples due to plant death and missed datapoints. To rule out thigmomorphogenesis, half of each plot for inbred lines was evaluated weekly, and the other half was evaluated once at the end of the growing season in 2021 and 2023 (**Figure S3**). For hybrids, measurements were taken from 3-9 symptomatic and 3- 9 adjacent symptomless plants per field. For both inbreds and hybrids, stalk diameter was measured at a height of 0.54 m (height of displacement) with a vernier caliper at every measurement.

### Destructive Quantification of Stalk Mechanics

3-point bend tests were performed on inbreds between 57-109 dap in 2024. Stalks were stripped of leaves and cut with pruning shears to a uniform length of 0.3 m, centered around the 0.54 m height. Stalks were measured within 1.5 hours of cutting and within the same time window as field-based measurements. This 1.5 hour window allows for minimal effects of moisture on the structural stiffness (K) and bending modulus (E) as maize stalk tissues have been shown to retain at least 70 % moisture within 4 hours after cutting <u>(Zhang et al. 2016; Zhang, et al. 2017;</u> <u>Sutherland et al. 2023)</u>. 3-point bend tests were performed using an Instron Universal Testing Stand Model 68SC-05 equipped with a 500 N load cell and an adjustable 3-point bend fixture (Instron Model 2810-400). The span length was set at 200 mm and samples were centered on lower anvils. Data was acquired with the Instron software (Bluehill 3.0). Digital calipers were used to measure the stalk diameter perpendicular (width of the stalk) and parallel (depth of the stalk) to the plane of displacement. Samples were preloaded to 0.2 N and displaced at a constant rate of 1.50 mm/s for 30 mm or until failure. Force-displacement data were exported from the Bluehill software and subset to include displacements of 0-2 mm. A linear regression was run with displacement as the independent variable and force as the dependent variable. The structural stiffness (*K*) was defined as the slope of the linear regression (N/mm). The second moment of area *(I*) was calculated using the equation for a solid cylinder (Hostetler et al., 2022; Niklas and Spatz, 2012):

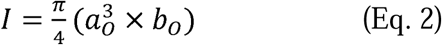

where n = 3.1415, a^3^_o_ is the radius parallel to the plane of displacement (mm), and b_O_ is the radius perpendicular to the plane of displacement (mm). The bending modulus (*E*) was calculated using:

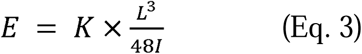

where *K* is the structural stiffness (N/mm), *L* is the fixture span length (200 mm for this study) and *I* is the second moment of area calculated in Eq. 2.

### Statistical analysis

All statistical analyses were performed in R ver. 4.4.0 (R Core Team, 2024) and all figures were generated with the ggplot2 ver. 3.5.1 (Wickham, 2016) and cowplot ver. 1.1.3 (Wilke, 2024) packages.

To model the stalk flexural stiffness, a logistic growth curve model was fit to the data (Bustos-Korts et al., 2019; Mu et al., 2022). Specifically, a nonlinear regression model (*nls* function) was used in combination with the *SSlogis* function where stalk flexural stiffness was the dependent variable and dap or GDD were the independent variable. The *nls* function fits a nonlinear regression model while the *SSlogis* function represents the logistic growth model. The logistic growth model can be explained by the following equation:

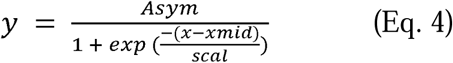

where y is the predicted stalk flexural stiffness, Asym is the maximum value of stalk flexural stiffness, xmid is the midpoint of the logistic curve, and scal is how fast stalk flexural stiffness changes around xmid. The *nls* function estimates the Asym, xmid, and scal of the data to minimize the sum of squared differences between the observed and predicted values. A logistic growth curve model was fit to each of the inbreds within each year and to each commercial hybrid genotype and disease state combination. While individual plant variability was observed, all data points were included in logistic growth models.

To determine if changes in stalk flexural stiffness are due to thigmomorphogenesis, plants that were measured repeatedly (weekly) were compared with those that were measured once with an analysis of variance (ANOVA) using the *agricolae* package ver. 1.3.7 (De Mendiburu, 2023). A one-way ANOVA was used within each genotype and year combination to compare the differences between groups, where the stalk flexural stiffness was the dependent variable and testing frequency was the independent variable. To determine if residuals were normally distributed, a Shapiro-Wilk test was used. If the residuals were not normally distributed, a Tukey’s Ladder of Powers from the *rcompanion* package ver. 2.4.36 (Mangiafico, 2024) was used to transform the data prior to running the ANOVA.

### Data availability

All raw data and R scripts used to process and analyze data are available at: https://github.com/EESparksLab/Ikiriko_Hostetler_et_al_2025. All code associated with the Pusher device is available at: https://github.com/EESparksLab/Pusher.

## RESULTS

### Stalk flexural stiffness has a biphasic trajectory

Stalk flexural stiffness was measured for three maize inbred and three commercial hybrid genotypes (**Figure 1A & B**). Inbred genotypes were measured across 2-3 years to capture environmental variability, whereas commercial hybrid genotypes were measured in one year but included both PRR symptomatic and symptomless plants. To define the overall trajectory of the stalk flexural stiffness, a logistic growth curve model was fit to each unique combination of year and genotype for inbreds, and genotype and disease state for hybrids. For all genotypes, the model demonstrates a biphasic trajectory composed of a linear increase followed by a sustained phase (**Figure 1C & D, Supplemental Table S1-S2, Figure S4**). The timing at which plants transitioned from the linear increase to sustained phase varied by genotype but was consistent within a genotype (occurring within 10 days).

**Figure 1.**
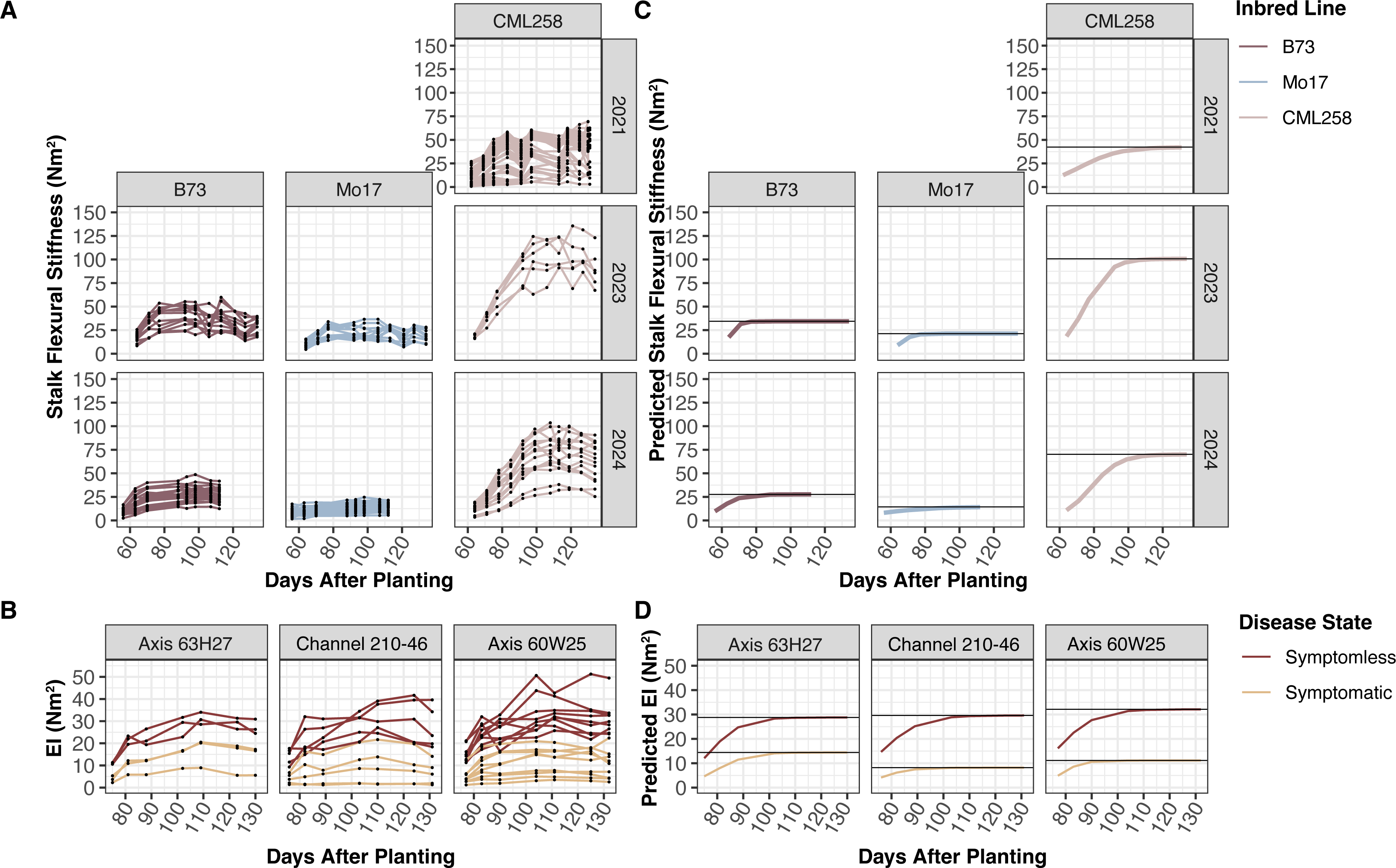
Stalk flexural stiffness follows a biphasic trajectory consisting of a linear increase and a sustained phase. Stalk flexural stiffness (*EI*) of **(A)** three inbred genotypes (B73, Mo17, and CML258) across multiple years and **(B)** three commercial hybrid genotypes with symptoms of Pythium Root Rot (yellow) and those that are symptomless (red). A logistic growth model was fit to **(C)** each genotype and year combination and **(D)** genotype and disease state combination and shows a consistent biphasic trajectory.

For inbred genotypes across environments, there was an altered the rate of increase, which changed the end-of-season stalk flexural stiffness. For CML258, the rate of increase was 121 Nm^2^/day (2021 environment), 182 Nm^2^/day (2023 environment) and 146 Nm^2^/day (2024 environment) (**Figure 1C**). The linear increase phase of B73 and Mo17 was likely also altered, but too abbreviated to determine the rate of change. In commercial hybrid fields, symptomatic plants that were marked for displaying early-season symptoms of PRR also had an altered rate of increase for stalk flexural stiffness (**Figure 1D**). For symptomless plants, the rate of increase was 44 Nm^2^/day, 54 Nm^2^/day, and 57 Nm^2^/day for Axis 63H27, Channel 210-46, and Axis 60W25 respectively. For symptomatic plants, the rate of increase was 22 Nm^2^/day, 11 Nm^2^/day, and 11 Nm^2^/day, respectively. While the rate of increase was impacted by environment and disease state, the trajectory of stalk flexural stiffness still followed a biphasic trajectory, and the timing of transition remained constant within a genotype.

### The timing of transition to the sustained phase is determined by physical time

We hypothesized that the timing of transition between phases was associated with plant growth stage. To test this hypothesis, the cumulative growing degree days (GDD) was used to normalize developmental progression across years (**Figures S1A**). The overall trajectory of the stalk flexural stiffness was again fit with a logistic growth curve model (**Figure S1B**). Between years there was a substantial offset of the transition point by 500-600 GDD within a genotype (**Figure S1B**). As expected, anthesis was observed at the same GDD across years (**Figures S1A**). Thus, these data show that the transition is not associated with the plant growth stage and is genetically encoded to occur relative to physical time.

### Changes in stalk flexural stiffness are retained after normalizing for diameter

The flexural stiffness measurements from the field are based on Euler-Bernoulli beam theory for a cantilever beam. This theory defines the flexural stiffness (sometimes called flexural rigidity) of a beam with a fixed base based on the height of deflection. The flexural stiffness measurement is determined by both the geometry of the stalk and the material properties of the stalk (**Figure S5A**). Separating the contribution of *E* and *I* requires quantification of the material properties (*E*), which can then be used to calculate the contribution of *I*. However, plants are composite materials and do not have a defined *E*, which limits the ability to separate these two contributing factors. To provide an initial indication of the contribution of geometry, stalk diameter was measured at each timepoint of testing. There was minimal change in stalk diameter within a genotype across the testing period (**Figure 2A** & **B**). Consistent with this data, when stalk flexural stiffness was normalized by diameter, the trajectory remained the same (**Figure 2C** & **D**), indicating that the trajectory of stalk flexural stiffness is unlikely to be driven by geometry.

**Figure 2.**
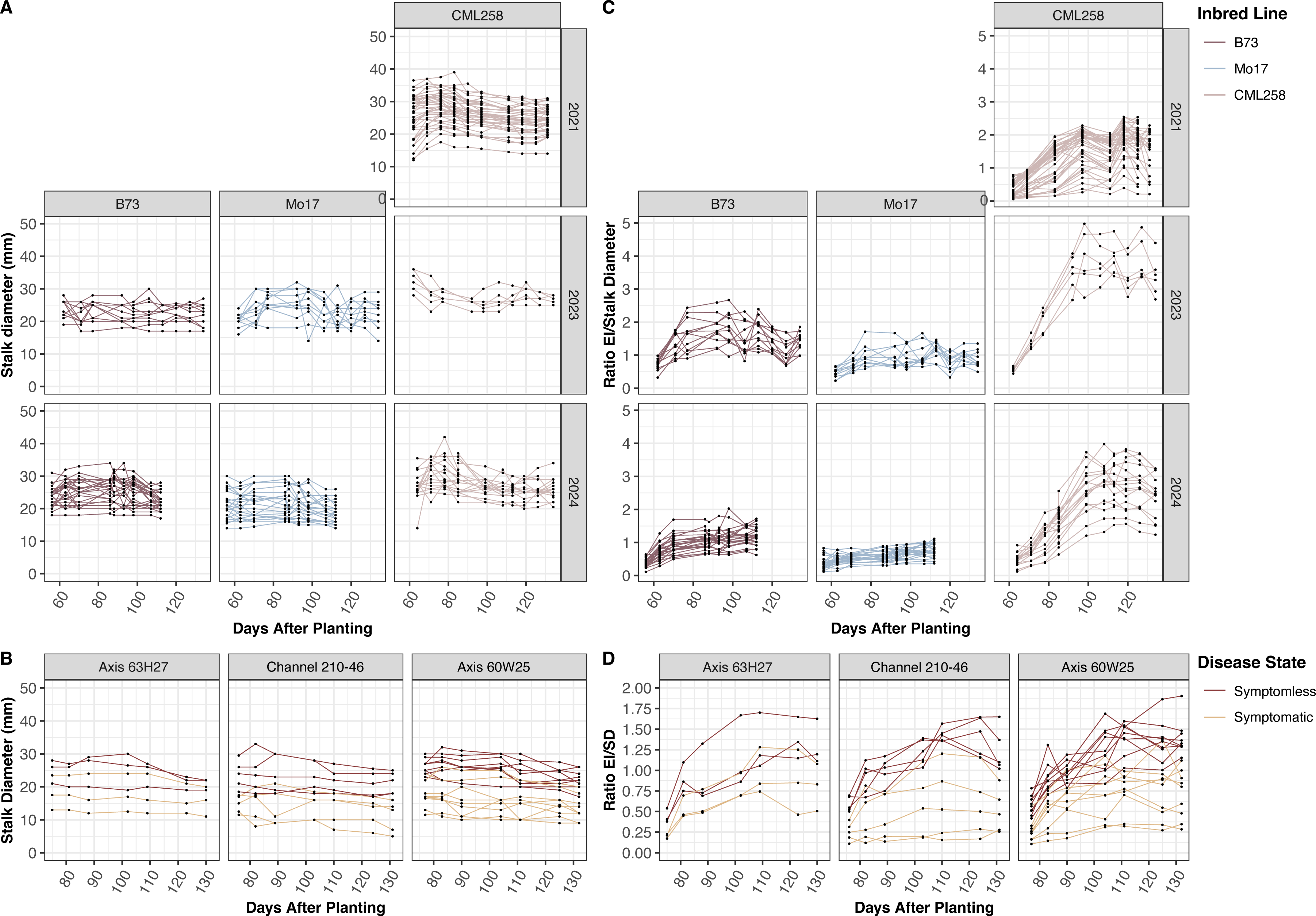
Stalk geometry is consistent while stalk flexural stiffness increases. Stalk diameter changes minimally over time for **(A)** inbred genotypes within years and **(B)** commercial hybrid genotypes with symptoms of Pythium Root Rot (yellow) and those that are symptomless (red). **(C** & **D)** The ratio of stalk flexural stiffness (*EI*) to stalk diameter (SD) maintains the defined biphasic trajectory of stalk flexural stiffness alone.

### Changes in stalk mechanics are driven by material properties

As highlighted above, field-based measurements are limited in their ability to separate the material properties (*E*) from the geometry (*I*) with the compound measurement of EI. Therefore, destructive 3-point bend tests were used to complement the field analyses. A 3-point bend test measures the mechanics at the mid-point of a section of the stalk supported by two lower beams, but with both ends of the stalk free (**Figure S5B**). The 3-point bend test is an inherently different measure then the field-base measures and therefore cannot be directly compared. However, the structural stiffness (*K*) from the 3-point bend tests followed a comparable biphasic trajectory as stalk flexural stiffness (EI) from the field (**Figure 3A**). Thus, demonstrating a consistent trajectory was identified using both types of measurements. Consistent with a lack of change in geometry in the field, the second moment of area (*I*) was also unchanged across the testing period (*p* > 0.05, **Figure S6**), emphasizing the consistency in geometric contribution. Indeed, the bending modulus (*E*), which removes the effect of geometry, retains a biphasic trajectory (**Figure 3B**). Collectively, these data demonstrate that the trajectory of stalk mechanics is driven by the material properties and not geometry.

**Figure 3.**
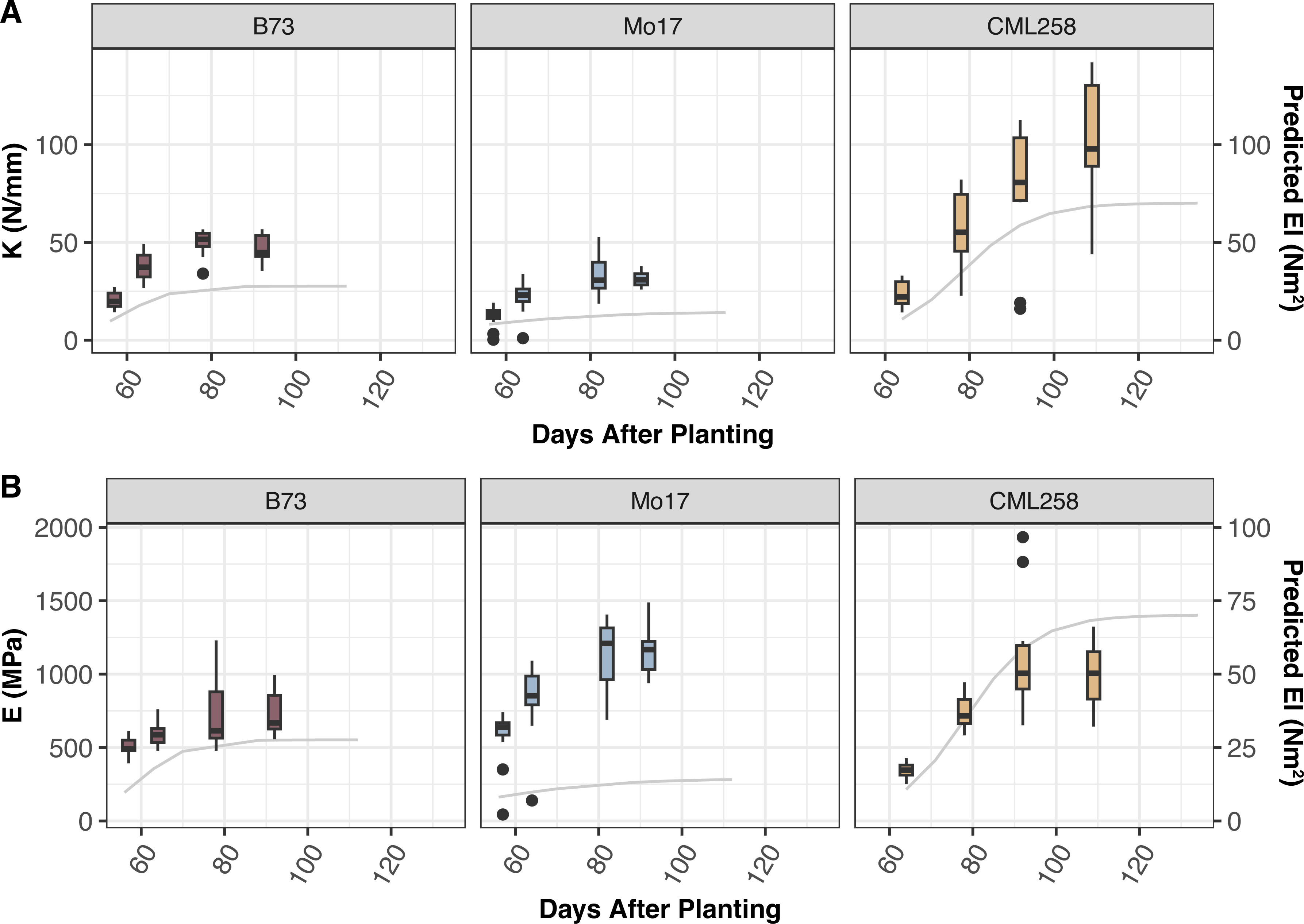
The trajectory of stalk flexural stiffness is driven by changes in the bending modulus. **(A)** Structural stiffness (*K*; boxplots) of inbred genotypes as determined by destructive 3-point bend tests follows the same trajectory as field-based non-destructive stalk flexural stiffness (*EI*; line graph). **(B)** The bending modulus (*E*; boxplots) from destructive tests also follows the same trajectory. Thus, demonstrating that material properties primarily drive the trajectory of stalk flexural stiffness.

## DISCUSSION

The trajectory of stalk flexural stiffness leading to variable end-of-season mechanics was previously undefined in maize. This study defined the trajectory of both research inbred and commercial hybrid maize genotypes as containing two phases – a linear increase followed by a sustained phase. This biphasic trajectory was also consistent across environments for inbred genotypes and in hybrid plants symptomatic for PRR. The rate of increase was altered by changes in either environment or disease state (**Figure 4A**). In contrast, the timing of transition to the sustained phase varied between genotypes, but was consistent within a genotype (**Figure 4B**) . These data demonstrate that the timing of transition to the sustained phase is genetically determined, while the rate of increase can be modulated within a genotype by environmental or biotic factors.

**Figure 4.**
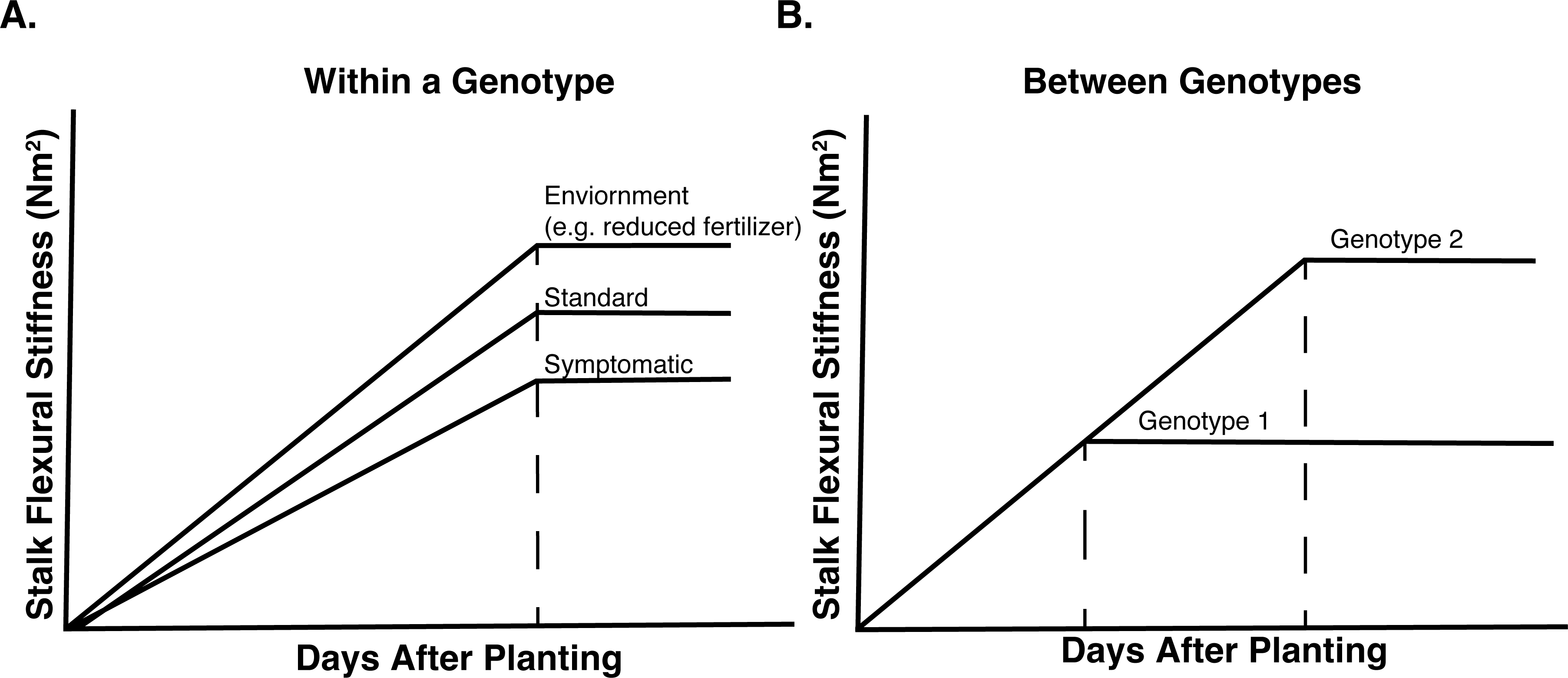
The biphasic trajectory of maize stalk flexural stiffness can be modulated by external and internal factors. **(A)** Within a genotype the rate of linear increase was changed by external factors such as fertilizer availability and disease state. **(B)** In contrast, the timing of transition from the linear to sustained phase was different between genotypes and thus likely controlled by internal genetic factors.

Results from destructive 3-point bend tests show that the bending modulus follows the same biphasic trajectory as non-destructive measures of stalk flexural stiffness. This indicates that changes in stalk mechanics are driven by material properties and not geometry. Previous end-of-season studies in maize hybrids suggested that stalk mechanics are primarily defined by geometry rather than material properties (Robertson et al., 2017; Stubbs et al., 2022a). However, there are substantial differences between these studies and the one presented here. Specifically, this study uses fresh stalk samples measured within 1.5 hours of cutting. In contrast, the previous studies analyzed senesced stalks and used dryers to reduce stalk moisture content, which will alter both stalk geometry and mechanics. Similar to the results presented in this study, differences in stalk mechanics in wheat were also attributed to material properties rather than geometry (Crook et al., 1994).

The bending modulus (i.e. material properties) is determined by the arrangement of cells within the stalk and the composition and organization of the cell wall. In monocots, such as maize, the absence of secondary growth rules out the influence of cell arrangement. Therefore, the most likely explanation for the changes in the bending modulus is the composition and organization of the cell wall. Previous studies have shown differences in stalk mechanics are driven by modifications to cell wall composition <u>(Ahmad et al., 2023; Zhang et al., 2017, 2014)</u>, which directly influence tissue (material) properties. For instance, there are significantly higher concentrations of total cell wall material in pith tissue of maize inbred genotypes that are resistant to corn borer larvae attack compared to susceptible genotypes <u>(Barros-Rios et al. 2011)</u>. Further, increased deposition of soluble sugars in the cell wall, and sclerenchyma tissue thickness have been linked to higher stalk flexibility and reduced fracture <u>(Wang et al. 2020)</u>. These findings are consistent with results in this study, where the plants from 2023 have a steeper increase phase. We hypothesize that the reduced fertilizer application in 2023 due to a lack of side-dressing altered cell wall material composition, resulting in higher stalk flexural stiffness. Future studies using defined fertilizer application strategies within a year, and quantifying the cell wall changes would help test this hypothesis.

The inbred genotypes that were assessed in this study included two temperate inbred genotypes, B73 (Iowa, USA) and Mo17 (Missouri, USA), and one tropical inbred genotype, CML258 (Mexico). The temperate inbreds are adapted to a longer photoperiod and were grown at a latitude close to their adapted region, and the tropical inbred was adapted to a shorter photoperiod and growing outside its adapted region. Tropical inbreds are photoperiod sensitive with increasing photoperiod leading to delayed flowering time (Bonhomme, Derieux, and Edmeades 1994). The timing of transition for B73 and Mo17 was quite similar, whereas the transition for CML258 was more delayed. Therefore, we hypothesized that the timing of transition would be linked to developmental stage or more specifically, flowering time. However, when assessing both growing degree days (general comparison of developmental stages) and timing of anthesis (flowering time), the timing of transition within an inbred did not overlap with flowering time (**Figure S1B**). We further explored the idea of cumulative solar radiation. Again, the timing of transition to the sustained phase did not overlap with a set cumulative solar radiation.

It remains unclear what signal plants are perceiving to allow them to monitor the physical passing of time, as opposed to integrated environmental signals regulating plant development. One possibility is the diurnal temperature range, which has been shown to alter vegetative growth and physiology (Sunoj et al. 2016). In our study, the average diurnal temperature range across all years is relatively consistent, varying within 1 (19.3 for 2021; 19.7 for 2023; 18.6 for 2024). Therefore, it is possible that plants can sense the diurnal temperature range, but additional studies are required to test this hypothesis. There may additionally be a photoperiod sensitivity mechanism that is independent of flowering time. Specifically, photoperiod determines diel rhythms (light-dark cycle), which in turn entrain the circadian clock. Among the many pathways that are altered by diel and circadian rhythms, cell wall components have been shown to be regulated in sugar cane grown under long or short day conditions (Alves et al. 2019). While this mechanism has not been determined for maize, we propose that this hypothesis is most consistent with our results. Further studies aimed at quantifying the cell wall responses in these lines will be important to define the genetic basis of transition timing.

Overall, these results show that the trajectory of maize stalk mechanics is characterized by a linear increase phase and a sustained phase. The rate and magnitude of these phases can be modulated by external factors such as nutrient application and disease, as well as internal factors such as genotype. While both geometry and material properties contribute to the trajectory of stalk mechanics, the data presented here show that stalk material properties drive the trajectory.

## Supplemental Data

**Figure S1. The trajectory of stalk flexural stiffness is not related to growing degree days. (A)** GDD accumulation for inbred genotypes in each year of analysis was used as a proxy for the plant growth stage. The points show the times when stalk flexural stiffness was quantified in the field. Horizontal black lines indicate the flowering date for CML258, B73, and Mo17 at the Newark, DE field site. **(B)** Logistic growth curve models of stalk flexural stiffness (*EI*) relative to GDD for each inbred genotype do not transition between linear increase and sustained phase at the same stage. The vertical black line within each panel indicates the flowering time (in GDD) for the respective genotype. The horizontal black lines indicate the ‘Asym’ value extracted from the logistic growth model and is defined as the maximum value of stalk flexural stiffness. The red star illustrates the timing of transition to the sustained phase.

**Figure S2. Overview of the Pusher device used to measure stalk flexural stiffness. (A)** The physical body of the Pusher device is composed of a (1) frame, (2) powertrain, and (3) load cell. **(B)** The internal electronics are run via an Arduino microcontroller. The microcontroller connects to a motor controller, load cell, and Bluetooth module. Dotted lines indicate a wireless connection, whereas solid lines indicate a hardwired connection between components.

**Figure S3. Changes in stalk flexural stiffness are not due to thigmomorphogenesis.** Within a plot, half the plants were tested weekly and the other half were tested once at the end of the season. A one-way ANOVA was conducted within each genotype and year and showed that testing frequency did not impact the stalk flexural stiffness (*p* > 0.05).

**Figure S4. Stalk flexural stiffness follows a biphasic trajectory consisting of a linear increase and a sustained phase.** A logistic growth model of stalk flexural stiffness (*EI*) was fit to **(A)** three inbred genotypes (B73, Mo17, and CML258) across multiple years and **(B)** three commercial hybrid genotypes with symptoms of Pythium Root Rot (yellow) and those that are symptomless (red). The shaded area surrounding the model is a confidence interval (95% confidence), which was determined via bootstrapping.

**Figure S5. Complimentary approaches to assess stalk mechanics. (A)** The Pusher measures the flexural stiffness (*EI*, Nm^2^) of stalk based on Euler Bernoulli beam theory. This theory considers the height of the measurement and assumes the beam has a rigid attachment at the base. (B) The 3-point bending approach measures the structural stiffness (*K,* N/m) at the midpoint of a stalk section that is supported by two lower anvils with both ends of the stalk section free.

**Figure S6. The second moment of area (I) is constant.** The second moment of area (*I*) was calculated for each stalk subject to 3-point bend tests. There were no differences in *I* across testing time points within a genotype (*p* > 0.05).

**Table S1: Logistic growth curve model metrics for maize inbred lines in 2021, 2023, and 2024.**

**Table S2: Logistic growth curve model metrics for maize commercial hybrid genotypes.**

**Video S1. Pusher device during testing.** Video showing the Pusher device with a load cell placed in contact with the stalk, slowly extending and retracting during testing.

## Acknowledgements

We gratefully acknowledge members of the Sparks lab (Jingjing Tong, Thanduanlung Kamei, and Emilia Pierce) for comments on the manuscript, UD Engineering Senior Design for initial Pusher prototypes, Sebastian Torres and Doris Cao for Pusher software modifications, Stephen Smith, Dave Griffin, Het Patel, Sebastian Torres, Joe Cristiano, Safiyya Haider, Amos Nyabuti, Austin Jensen, Kishan Biradar, and Lee Bowman for help with data collection, and Maddle Henrickson for flagging symptomatic PRR plants. Special thanks goes to RC Willin and Willin Farm LLC for allowing on-farm research and partnering for this work. This work is supported by Delaware Biotechnology Institute (DBI) Delaware Bioscience Center for Advanced Technology (CAT) and Applied Research Collaboration (ARC) Award #12A00448 to AKB; the National Science Foundation (NSF) Division of Civil, Mechanical and Manufacturing Innovation (CMMI) Program in Biomechanics and Mechanobiology (BMMB) Grant #2040346 to EES; the Pests and Beneficial Species in Agricultural Production Systems project award no. 2023-67013-39165 awarded to AKB and EES, and the Physiology of Agricultural Plants Postdoctoral Fellowship project award no. 2022-67012-36840 awarded to ANH, from the U.S. Department of Agriculture’s National Institute of Food and Agriculture.

## Abbreviations

Dap: (days after planting)
GDD: (Growing Degree Day)
PRR: (Pythium Root Rot)

