## Supplementary figures and images for "A biphasic trajectory for maize stalk mechanics shaped by genetic, environmental, and biotic factors"

### Figure S1

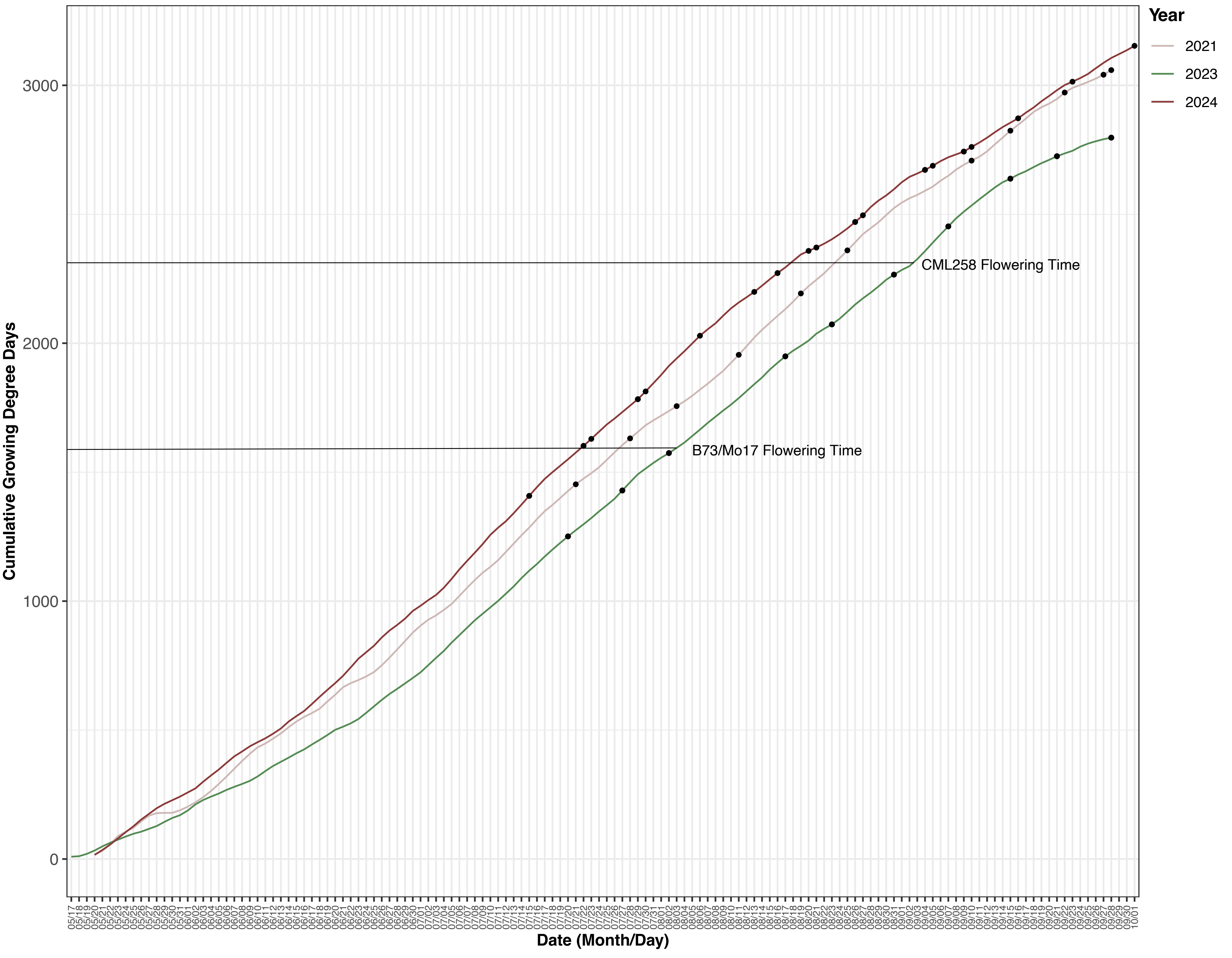

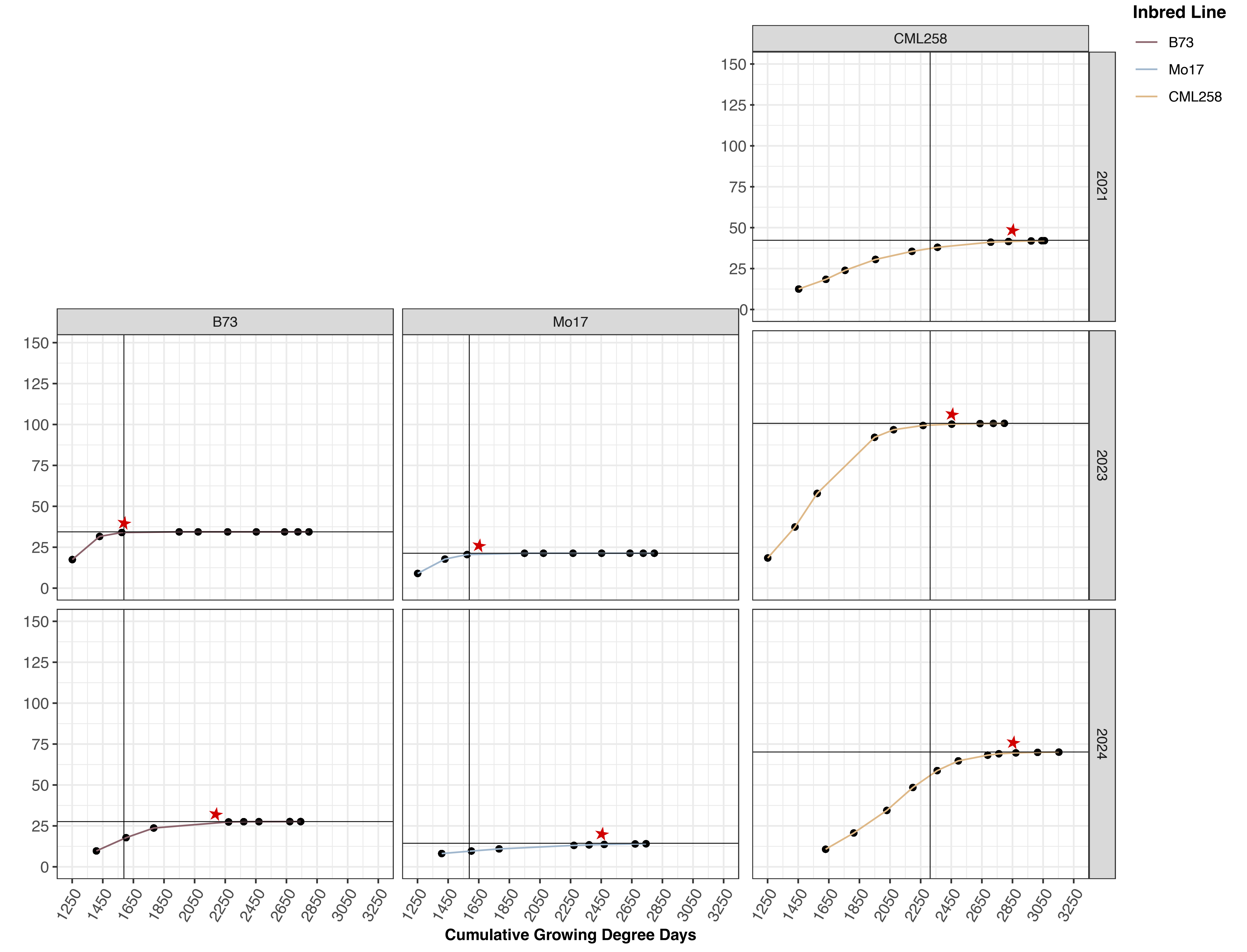

### Figure S2

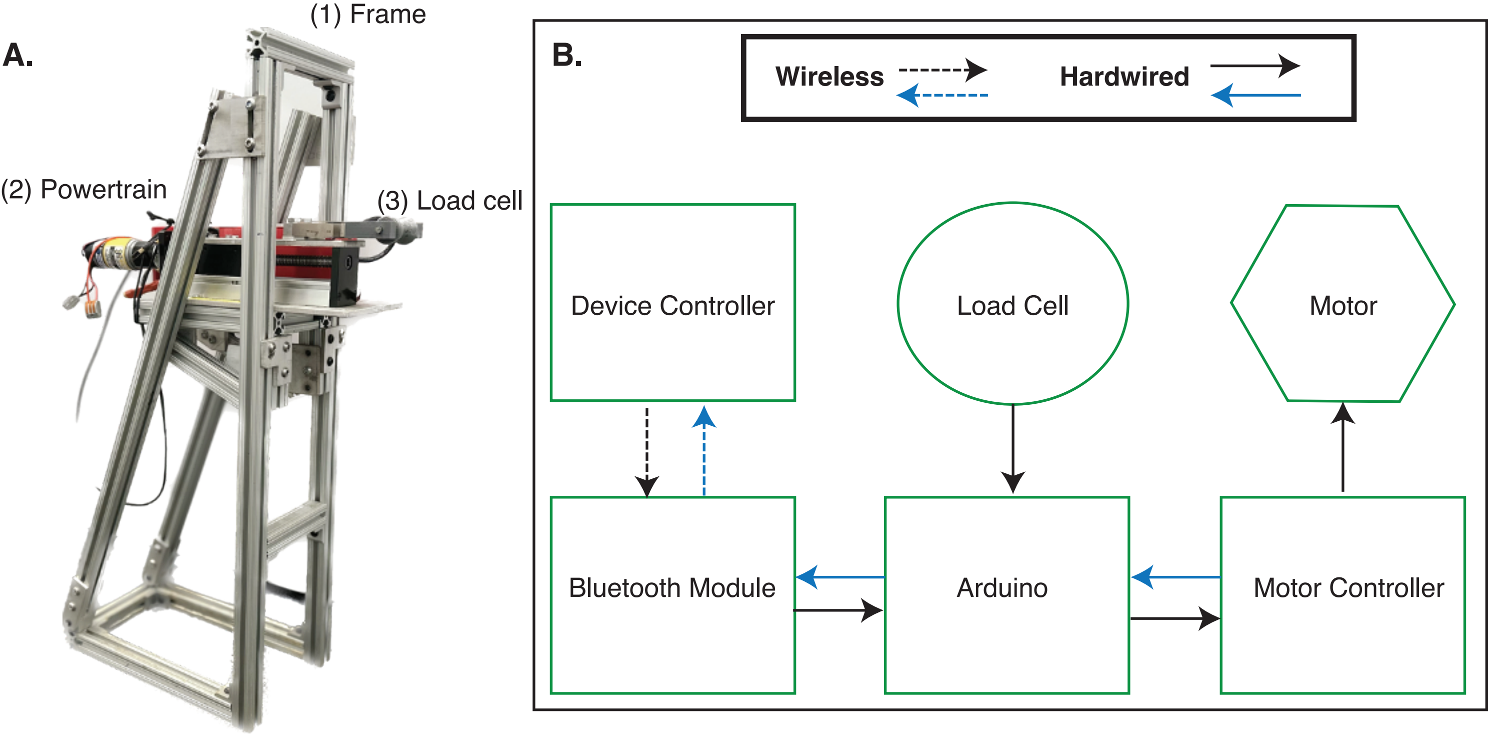

### Figure S3

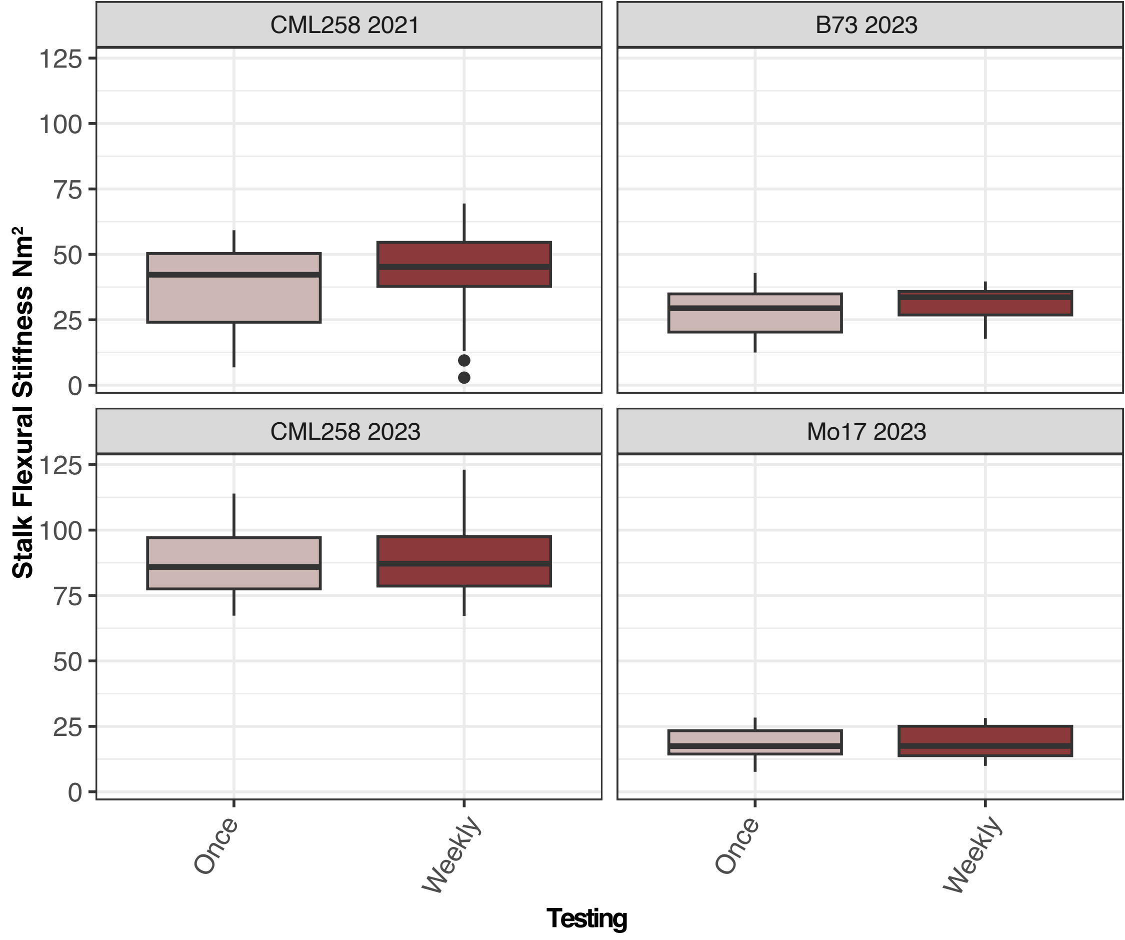

### Figure S4

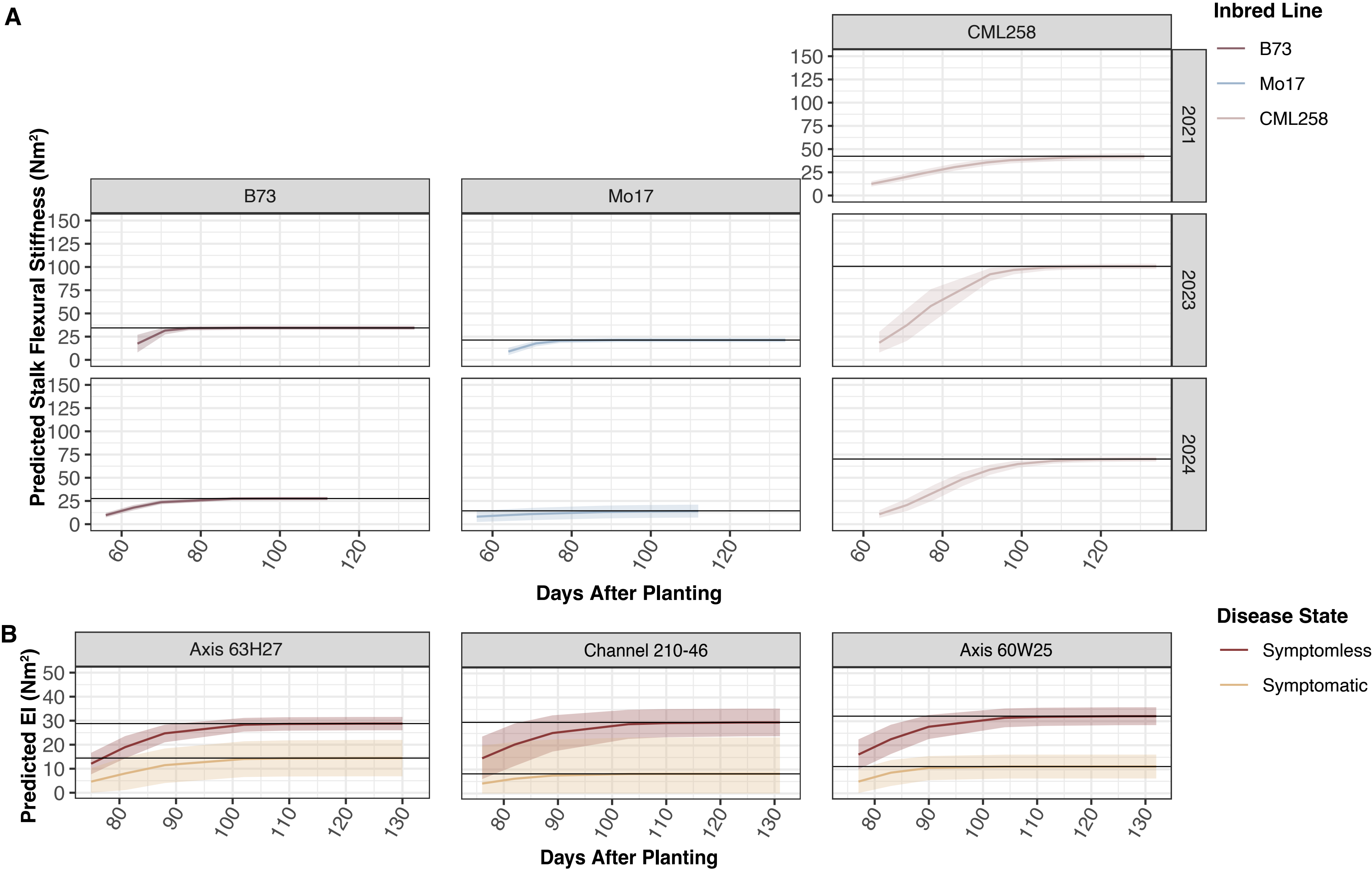

### Figure S5

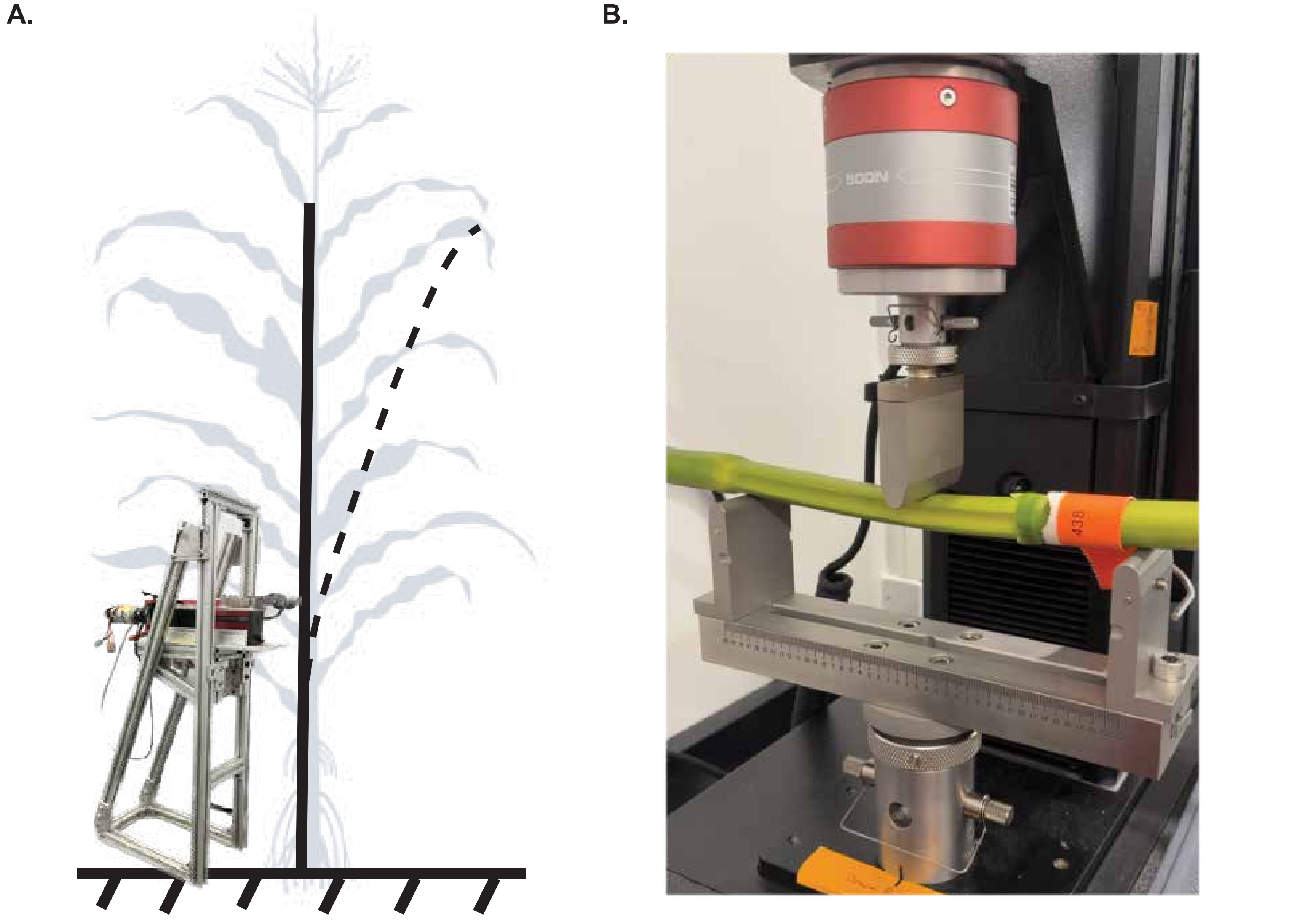

### Figure S6

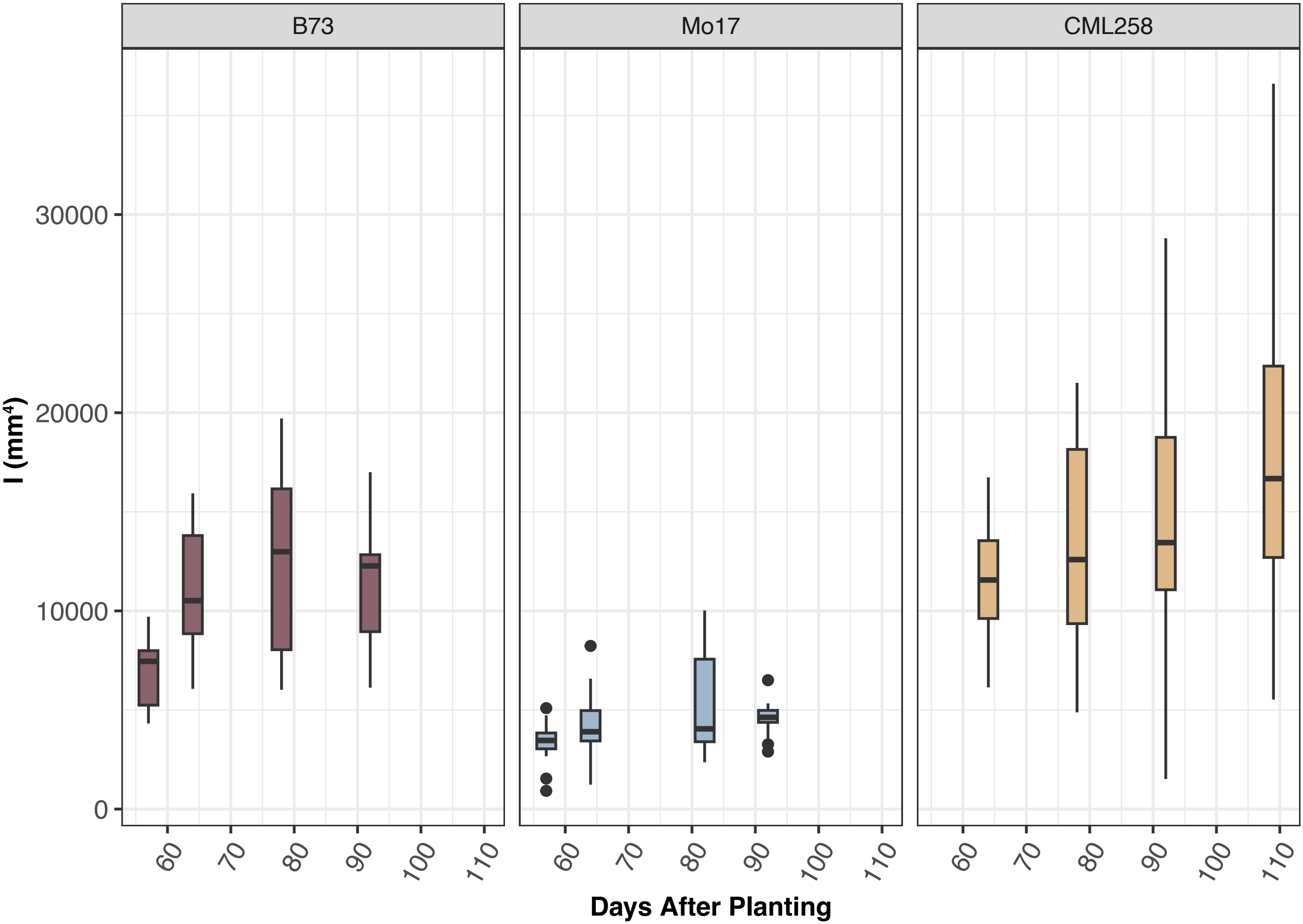
